# Diversity begets diversity under microbial niche construction

**DOI:** 10.1101/2022.02.13.480281

**Authors:** Sylvie Estrela, Juan Diaz-Colunga, Jean C.C. Vila, Alicia Sanchez-Gorostiaga, Alvaro Sanchez

## Abstract

Microbial interactions are expected to modulate microbial diversity, but whether they inhibit or stimulate further diversity in complex microbial communities, and how, remains poorly understood. By assembling microbial communities in multiple identical habitats with a single limiting nutrient, here we provide direct evidence for the role of microbial niche construction and cross-feeding in driving a positive relationship between community diversity and focal lineage diversity in microbial communities. Combining these experiments with simulations, we show that this positive relationship is not inevitable but depends on the underlying metabolic structure of by-product secretions and uptake between different taxonomic levels.

## Introduction

Microbial diversity is a critical determinant of microbiome functioning and ecosystem health. The effects and implications of microbiome diversity on the services and functions that microbiomes provide are often complex and can be context dependent [1, 2]. For instance, loss of microbial diversity in the human gut has been linked to intestinal dysbiosis and digestive diseases [3, 4], while at the same time increasingly diverse vaginal microbiomes have been associated with dysbiosis and infection [5]. Well-controlled manipulative experiments have found similarly complex responses of ecosystem function to biodiversity. Studies in laboratory fly populations reported that increasing microbiome diversity generally lowered the lifespan of these organisms [6], while similar manipulative studies in plant microbiomes found that increasing functional diversity protected the host against pathogens [7]. In model biorefinery communities, increasing biodiversity has a non-monotonic effect on the amount of biofuel produced [8].

These examples illustrate the central role that biodiversity plays in microbiome function, and motivate ongoing efforts to understand the ecological mechanisms that shape it. Microbiome diversity is governed by multiple environmental factors including niche availability (number and type of resources supplied externally), exposure to stressors, the frequency and intensity of disturbances, dispersal rates, habitat area, and the degree of spatial structure [9–14] (**Fig. 1**). In addition to these external factors, biodiversity may also be modulated by diversity itself, through the effect of species interactions in the exclusion and recruitment of additional taxa. How can biodiversity feed back upon itself? Two alternative hypotheses have been proposed [15]: one model, known as niche filling (in essence, negative feedback control), suggests that high biodiversity limits further increases in diversity due to increased competition for shared, limiting resources. A second alternative model, known as the ‘diversity begets diversity’ (DBD) hypothesis (i.e., positive feedback control), suggests that more diversity creates more ecological niches, thus promoting further diversification (**Fig. 1**). These hypotheses have been investigated in natural populations of plants and animals, often with contrasting outcomes [16–18]. In microorganisms, studies have primarily focused on laboratory experiments studying the evolutionary diversification of a single focal strain in small microcosmos (e.g., [19–21]).

**Figure 1.**
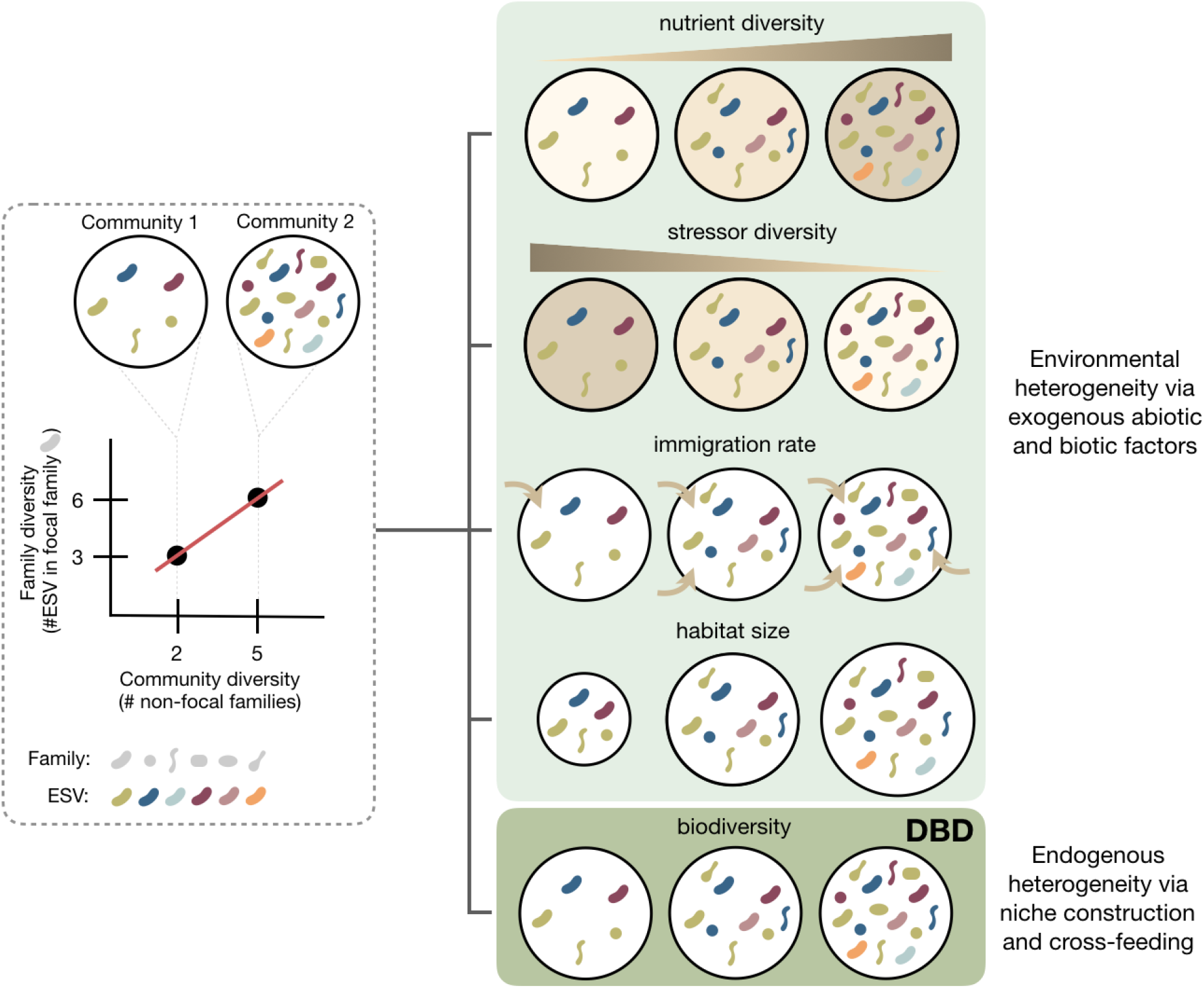
Drivers of microbial diversity in microbial communities. Microbial diversity may be shaped by both exogenous (environmental) and endogenous (microbial interactions) factors. We illustrate scenarios where increasing resource diversity, immigration rate, and habitat size all lead to an increase in diversity whereas increasing environmental stressors reduces diversity. Microbial interactions may similarly influence biodiversity, for instance through metabolic niche construction (i.e., secretion of metabolic byproducts) and cross-feeding among microbial taxa. We depict a scenario where as community diversity increases, the number of metabolites secreted-- and novel niches-- may also increase, creating further opportunities for cross-feeding and more diversity. In this scenario, diversity begets diversity (DBD), as diversity itself is a driving force for further diversification. Together, this illustrates alternative mechanisms by which a positive diversity slope (i.e., a positive correlation between community diversity and focal taxa diversity) may be observed across different environments.

The main difference between both hypotheses is that the DBD hypothesis predicts a positive correlation between diversity in focal lineages and existing biodiversity, whereas the niche filling hypothesis predicts a negative correlation (**Fig. 1**) [22]. While these two models may appear to be incompatible, they can actually describe different regimes of diversity: the DBD may apply at low diversity, whereas the niche filling hypothesis may manifest itself at higher species richness [22, 23]. Using published microbiome data from the Earth Microbiome Project (EMP) [24], Madi et al (2020) found evidence for a strong positive correlation between diversity in focal lineages and overall community diversity in less diverse (often host-associated) microbiomes like the gut microbiome, but often weaker correlation in more diverse microbiomes like in the soil, yet the underlying mechanisms behind these patterns remain unknown. Niche construction via metabolic cross-feeding has been proposed as a potential explanatory mechanism for DBD [22, 23], a point that is consistent with genome-scale metabolic modeling [23]. Yet, a positive correlation between community diversity and focal lineage diversity can in principle be caused by a variety of alternative mechanisms and not necessarily due to ‘diversity begetting diversity’ (**Fig. 1**). For instance, habitats may differ in the amount of a limiting resource they possess, such as space or nutrients, differ in residence times, or in the strength of migration (**Fig. 1**). These differences can lead to a positive correlation between community diversity and diversity in focal lineages, which can emerge from randomly sampling individuals into habitats with different population sizes (Supplementary Text; **Fig. S1**). Disentangling these and other potential mechanisms from the DBD hypothesis is very challenging in natural habitats, due to the inherent complexity and heterogeneity of natural microbial ecosystems and the large number of unknown factors that may be governing community composition in the wild.

To unambiguously demonstrate the effect of niche construction, and therefore show that diversity can pull itself by its figurative ‘bootstraps’, one would need to set up assembly experiments where diversity is the only (or at least dominant) variable between habitats. Multi-replicated enrichment communities, which consist of natural microbiomes cultivated in multiple identical, replicate habitats under well-defined growth medium and well-controlled conditions, are an ideal microbial system to test this hypothesis. Habitats may be inoculated from the same species pool and we can set them up to be exactly identical to one another except for the niches created by the species within, therefore removing the confounding effect of the (often unknown) sources of between-habitat variation that are inherent of natural habitats [25]. Enrichment communities can also be assembled in replicate habitats containing a single growth-limiting nutrient without immigration (besides the initial inoculation), and they lead to stable multi-species communities (5-36 ESVs) [10, 26, 27]. If we supply a single externally supplied limiting nutrient, most of the coexisting taxa will be inevitably supported by cross-feeding interactions among microbes in those communities [26, 27]. Further, even in identical habitats colonized from the same inoculum, communities tend to adopt alternative stable states, and thus diversity can differ spontaneously among replicate communities [26–28], allowing us to test the presence of a positive feedback in biodiversity without having to externally manipulate it ourselves.

Using multi-replicated enrichment communities assembled in a single limiting carbon source, here we set out to investigate whether DBD occurs in microbial communities where most of the available resources (niches) are generated by the microbes themselves, rather than being externally supplied. Our experimental setup allows us to test the DBD hypothesis under conditions where the only variation between habitats is endogenously created by the microbes living therein.

## Results and Discussion

### Diversity begets diversity in microbial communities assembled in a single nutrient

As discussed above, in order to provide a direct test of the DBD hypothesis we set out to design an experiment where communities would be assembled in well-controlled, externally identical habitats under a single supplied limiting nutrient so that cross-feeding would be required for any ecological interactions. Our experiments included 742 self-assembled enrichment communities in total, which were assembled in 24 different environments. Briefly, 8 different soil inocula were seeded into M9 minimal media supplemented with a single carbon source (24 different carbon sources in total, including 12 sugars and 12 organic acids) in 3-4 replicates each (Methods) (**Fig. 2**). Communities were passaged to fresh media (125 fold dilution) every 48h for 10 Transfers (Methods). The taxonomic composition of each community at Transfer 10 was assessed using 16S rRNA amplicon sequencing (Methods). To avoid sequencing depth biases in diversity estimation, all samples were rarefied to the same depth (10779 reads per sample).

**Figure 2.**
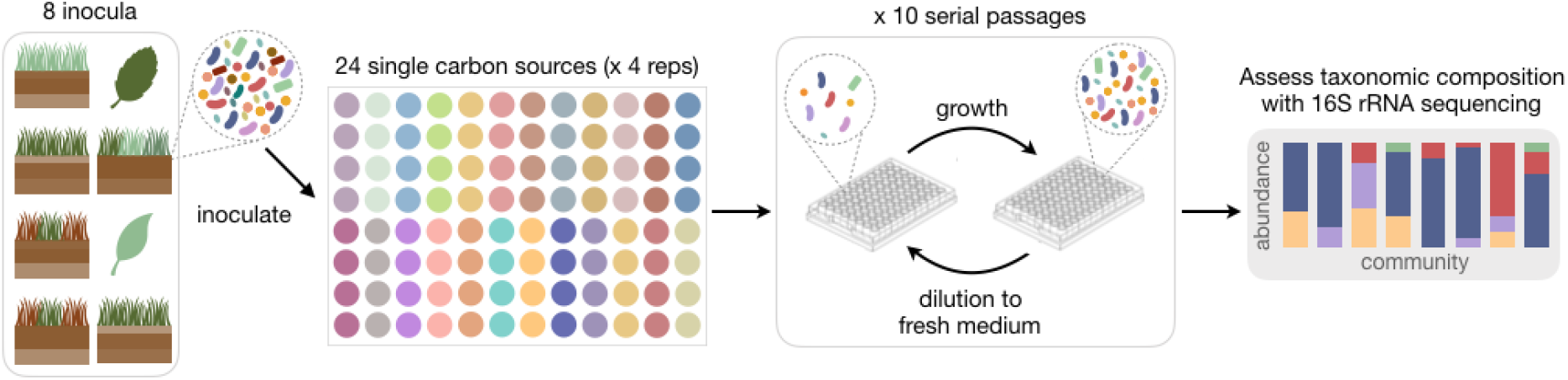
Experimental design of the microbial community assembly experiment. Microbial communities from 8 different environmental sources were inoculated into minimal medium supplemented with a single carbon source (24 carbon sources in total) in 3 or 4 replicates each (Methods). Communities were allowed to grow for 48h and then diluted to fresh medium (125x dilution). This growth-dilution cycle was repeated 10 times. At transfer 10, the communities were sequenced using 16S rRNA amplicon sequencing to assess the taxonomic composition (Methods).

To detect DBD in our communities, we followed a similar approach to the one used by Madi et al (2020): we studied how the number of Exact Sequence Variants (ESVs) belonging to a given focal family changes as the number of additional families present in the community increases. To quantify this dependence, we estimated the slope of the linear regression line, also referred to as the diversity slope [22]. A positive slope is consistent with diversity promoting diversity while a negative (or flat) slope is consistent with diversity limiting itself by niche filling. Here we specifically focus on the ratio between diversity at taxonomic level *i* and diversity at taxonomic level *i*+2 (i.e., ESV:Family), unlike Madi et al (2020) who calculated the diversity slopes between two consecutive taxonomic levels (e.g., ESV:Genus and Genus:Family). Primarily, this is because in previous work we have found that the family level of taxonomy provides a good proxy for ecological function under our conditions [27]. In addition, our communities generally have a smaller number of taxa, and therefore the ratio between diversity at taxonomic level *i* and diversity at taxonomic level *i*+1 can be too small for statistical significance. Finally, genus-level taxonomic assignment is generally less accurate and more variable [29]. Thus the ESV:Family diversity slope gives us both a greater and more accurate diversity range in our experimental communities.

We started by aggregating all habitats together and estimating the diversity slope of all communities for all families (a total of 360 ESVs spanning 46 families) and found a very weak positive diversity slope (m=0.02, R^2^=0.002, p<0.001, N=6007) (**Fig. 3**). Breaking it down by family reveals that a positive slope is observed for the four most abundant families across our communities: Enterobacteriaceae (m=0.08, R^2^=0.02, p<0.001, N=742), Pseudomonadaceae (m=0.36, R^2^=0.28, p<0.001, N=731), Moraxellaceae (m=0.04, R^2^=0.05, p<0.001, N=629), and Rhizobiaceae (m=0.07, R^2^=0.05, p<0.001, N=597) (**Fig. 3**).

**Figure 3.**
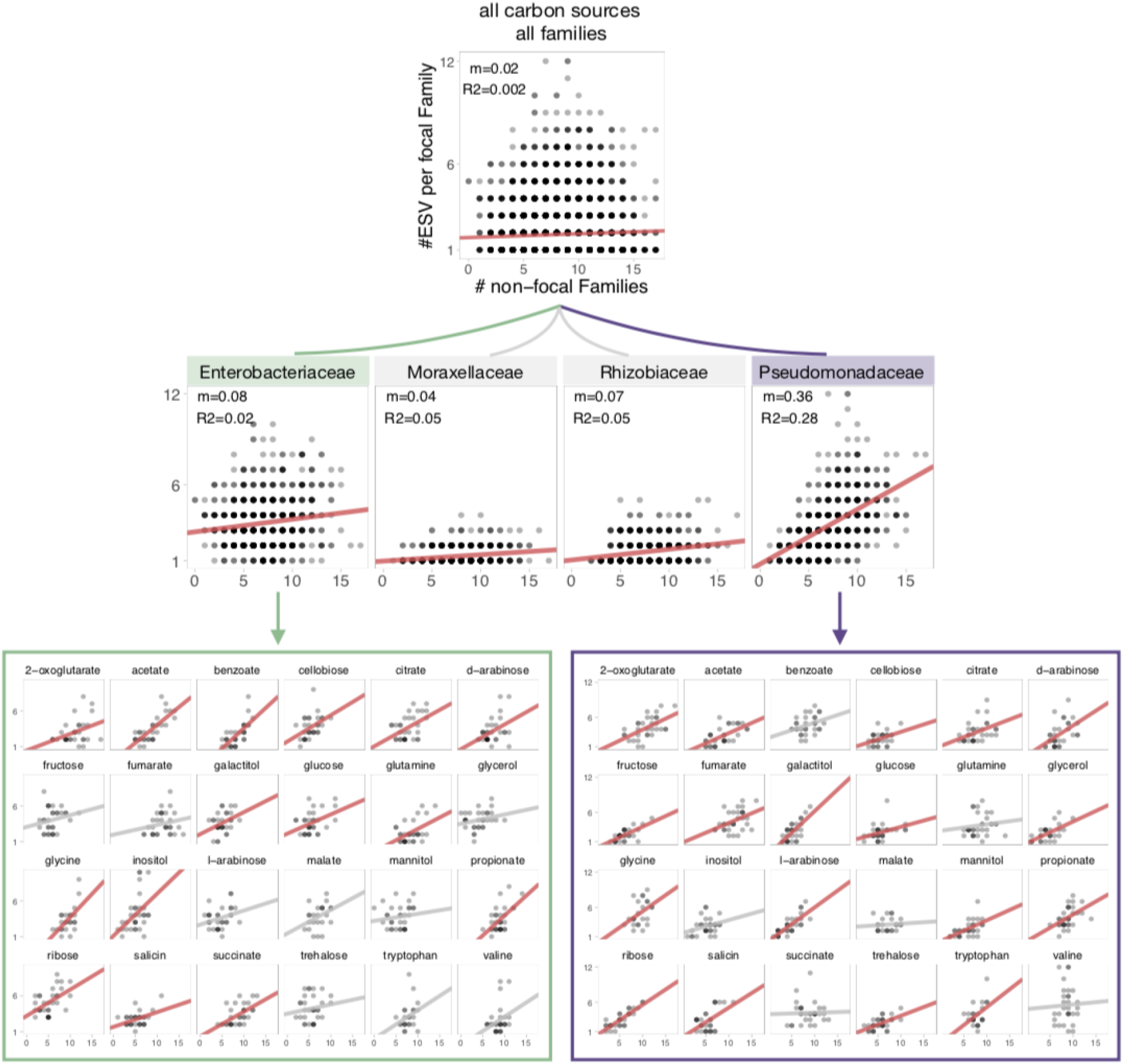
Communities self-assembled in a single carbon source exhibit a strong DBD pattern, but its magnitude may be masked by aggregating different habitats. All plots show the focal lineage diversity (number of ESVs per focal family) as a function of community diversity (number of non-focal families). For visual clarity, only the top panel has labeled axes. The plots show the diversity slope for all habitats and all families aggregated together (top row), all habitats aggregated by family (middle row), or separated by habitat and family (bottom row). Shown are the four most abundant families (Enterobacteriaceae, Pseudomonadaceae, Moraxellaceae, and Rhizobiaceae), which together represent ~88% of the total community abundance across all N=742 communities (E~47%, P~27%, M~8%, R~6%). The other families are not shown. Individual points are partially transparent, thus regions of the plot with higher density of points appear darker. The line represents the regression line (linear model). Significantly positive slopes (m) are shown in red and non-significant slopes are shown in gray (p>0.05).

The above result aggregates over different environments. The design of our experiment allows us to test the DBD hypothesis directly, by examining the diversity slope in a set of replicate habitats where the only variable that may be different between habitats is the different species interactions that may be taking place in each, i.e. diversity itself. To this end, we segregated communities by the limiting carbon source in their habitat, and plotted the number of ESVs in each focal family against the number of families in each replicate habitat. Consistent with the DBD hypothesis, we observed a positive slope for the two dominant families (Enterobacteriaceae and Pseudomonadaceae which consist of ~47%and ~27% of the total average abundance in the metacommunity, respectively) in each of the 24 environments, the majority of which are statistically significant (**Fig. 3**).

### Cross-feeding interactions drive the emergence of DBD

Next, we asked whether the observed positive slopes are expected to generically emerge in any community where the diversity is within the same range of our experiments. For this, we developed a null model of community assembly that considers no interactions across community members, the composition of a community is determined by which species in the inocula are passively selected by the environment (Supplementary Text). The model reveals that even a random sample of species from a highly diverse pool can exhibit a positive slope (with the effect being stronger for smaller sample sizes) (**Fig. S1**). Thus we asked whether the slopes observed in our self-assembled communities are simply caused by this sampling size effect, or on the other hand, our communities display stronger diversity slopes than what would be expected in the absence of interactions. For that, we considered a null model in which the entire collection of ESVs in our communities are randomly swapped across samples (Methods), thus breaking any potential interactions between species that result from community assembly. For both Enterobacteriaceae and Pseudomonadaceae, we find that close to half of the diversity slopes in the 24 carbon sources are greater than expected by chance (p<=0.05, N=500 randomizations; **Fig. S2A**). Similarly, the R^2^ observed is also generally stronger than expected by random chance, especially for Pseudomonadaceae (in 15 out of the 24 carbon sources, p<=0.05, N=500 randomizations; **Fig. S2B**).

A potential confounding factor still exists in our experimental design. We want species interactions to be the only difference across habitats. Yet, eight different inocula were used for each of the 24 different carbon sources (each replicated three or four times). In principle, one may conceive that the different initial diversity of each inocula could determine the final diversity of the replicate communities assembled from it. To test this hypothesis, we plotted the number of ESVs per focal family as a function of the number of non-focal families for each of the inocula, and calculated the regression (i.e., diversity) slope. Inconsistent with this hypothesis, we find no positive diversity slope for the inocula (m=-0.03, p>0.05, N=207; **Fig. S3A**). In addition, there is also no positive correlation between inoculum richness and the richness of the self-assembled communities both at the ESV and Family level (**Fig. S3B-C**). Together, this suggests that the positive slope we observe in the assembled communities cannot be explained by inocula diversity.

Our results suggest that interactions across community members, rather than either sampling effects or exogenous factors, drive the emergence of the strong positive diversity slopes that we observe in our experiments. Is this expected to be the case whenever community assembly is dominated by cross-feeding interactions across species? In other words, does cross-feeding inevitably lead to diversity begetting more diversity?

To examine this question, we turned to a Microbial Consumer Resource Model (MiCRM) which has been shown to effectively capture the dynamics of microbial communities in environments like the ones we used in our experiments (well-mixed medium with a single externally supplied limiting resource) [26, 30–32]. Our simulation procedure and a full description of the model are explained in detail in the Methods section. Briefly, we started by generating a library of resources and a pool of species (consumers) (Methods). Each consumer species is characterized by a unique vector of uptake rates of each of the resources. Resources are divided into classes, and consumer species are divided into families, such that every family of species displays preference (i.e., higher uptake rates) for a certain resource class (Methods). This reflects empirically observed differences in quantitative nutrient utilization traits at the family-level (e.g., Enterobacteriaceae’s preference for sugars and Pseudomonaceae’s preference for organic acids) [27]. In the MiCRM, species secrete metabolic byproducts into the environment. Which byproducts are secreted, and in what amounts, is encoded in a metabolic architecture (Methods) that can either be shared by all species across all families (if there exists a common ‘core’ metabolism, that is, every species secretes the same byproducts and in the same amounts when utilizing a same resource) or different across families of consumers (if species belonging to different families secrete different metabolites, and/or in different proportions, when consuming the same resource).

To determine whether a positive diversity slope is a generic emergent behavior of consumer-resource models, we ran 100 sets of simulations, assembling 100 communities in each one. Model parameters were randomized for each set (Methods), and communities were assembled so as to mimic our experimental design: we started by randomly sampling a subset of species from the pool, allowing them to grow to stationary phase, diluting their abundances by a factor of 1:100, and repeating the process for 20 cycles, at which point the composition of the communities was stable and did not change in subsequent generations (Methods). We then examined the relationship between the number of families and the number of species in the focal (most abundant) family (**Fig. 4A**).

**Figure 4.**
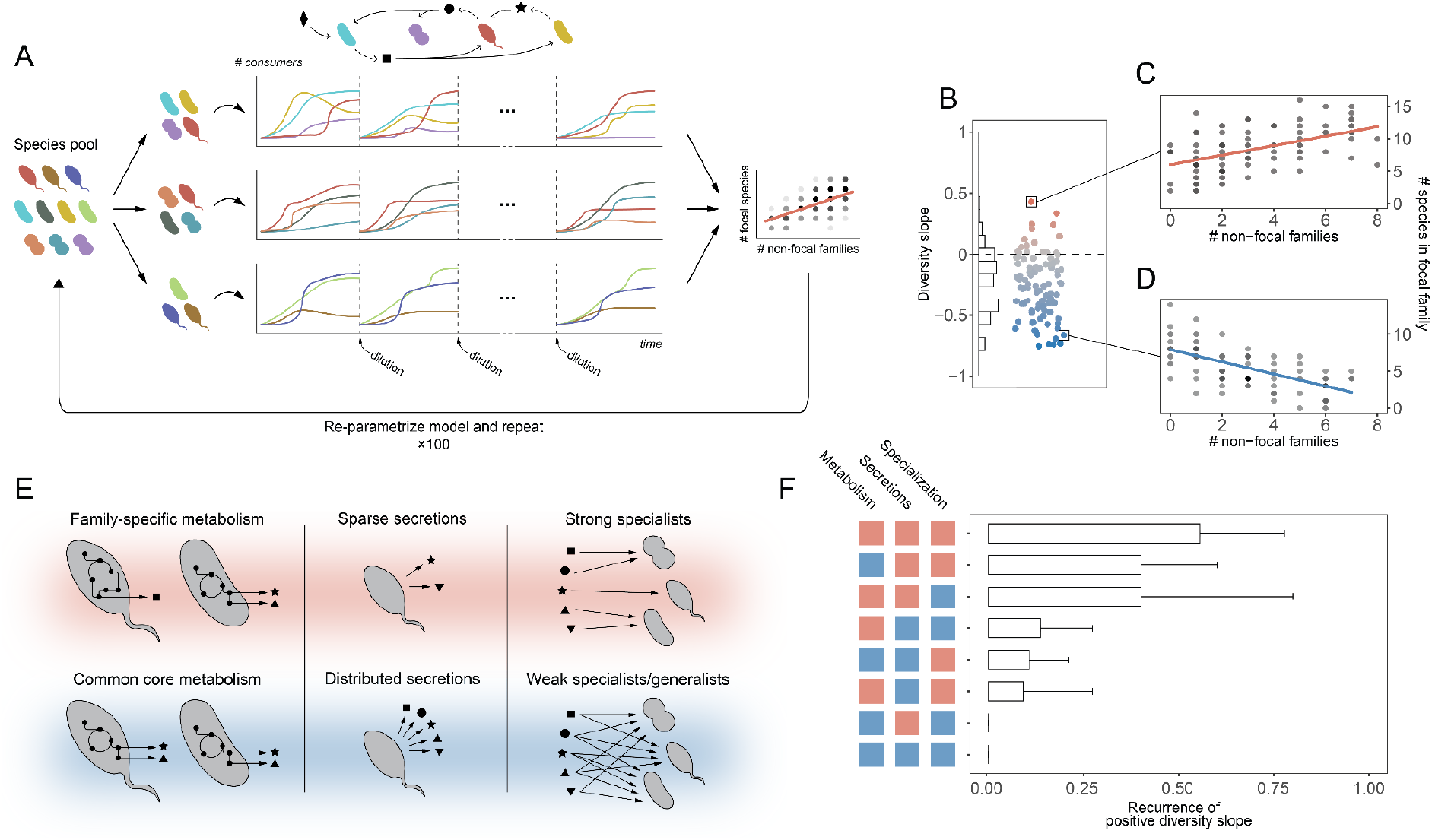
Emergence of DBD in a Microbial Consumer-Resource Model. (**A**) Simulations were carried out by first generating a library of species (consumers) divided into families. 100 communities were initialized by randomly sampling the species pool. Communities were stabilized by passaging them through 20 cycles of growth-dilution. The dynamics of the species between consecutive dilutions are determined by their interactions with one another via the secretion and uptake of metabolic byproducts, represented by the black shapes in the network. After stabilization, the number of families and the number of species in the focal family were quantified in each community and the diversity slope was obtained. The whole process was repeated 100 times, each with new model parameters (Methods). (**B-D**) Diversity slopes were often found to be negative, indicating that cross-feeding does not necessarily lead to DBD in the MiCRM. (**E**) Simulations were categorized according to three parameters: the family-specificity of the metabolic byproducts secreted by the consumers, the sparsity of those secretions, and the degree of specialization of the consumers as described in the main text. (**F**) Different parameter regimes lead to variable recurrence of DBD in the MiCRM. Colors match those in panel E: red for family-specific metabolic byproducts, sparse secretions, or strong specialization; blue for common metabolic byproducts, distributed secretions, or weak specialization. Error bars indicating 95% confidence intervals were computed by bootstrapping.

We found that diversity slopes were often negative or not significant (**Fig. 4B-D**), even though community assembly is dominated by cross-feeding interactions in all of our MiCRM simulations. This suggests that cross-feeding alone does not inevitably produce DBD, and that further underlying metabolic conditions may be required for a positive diversity slope to emerge. We reasoned that a positive diversity slope would emerge whenever the niches constructed by the families in the communities are different (at least in part) from family to family, and when these niches (at least some) can be exploited by additional members of the focal family. These conditions could be met by the MiCRM under some regimes and violated under others. For instance, if all species secrete the same or very similar metabolic byproducts into the environment regardless of the family they belong to, the niches constructed by different families would overlap substantially and adding a new family would not lead to more niches. Conversely, when secretions are family-specific ‘constructed niche overlap’ is reduced and adding new families can increase the number of niches (note that, in this context, ‘constructed niche overlap’ refers to the niches created by species through the secretion of metabolic byproducts, not to the overlap of species occupying similar niches). The degree of constructed niche overlap would also be minimized if byproduct secretions were distributed across a few different metabolites rather than across a wide variety of them. For a similar reason, if species were strong specialists of a small subset of resources (i.e., they exhibited a strong preference for them) instead of weak specialists or even generalists (i.e., they displayed weak or no preference for any resource class in particular), constructed niche overlap would also be reduced since the types of byproducts that can be secreted is contingent on the consumed resource and lower variation in consumed resources will naturally result in lower variation in secreted byproducts.

These factors can all be controlled by varying the parameters of the MiCRM (Methods). To test whether they modulate the emergence of a positive diversity slope in the model, we divided our set of simulations by whether byproduct secretions were common to all families or family-specific, whether secretions were evenly distributed across a wide variety of byproducts or sparsely distributed across only a few, and whether consumers were weak or strong specialists (**Fig. 4E**). For each subset of simulations, we quantified the frequency with which we observed a positive diversity slope. **Fig. 4F** shows that, in line with our reasoning, DBD tends to emerge substantially more often when byproduct secretions are family-specific and sparse, and when consumers are strong specialists. Previous community assembly experiments have shown that both nutrient consumption rates and secretions are quantitatively conserved at the family-level [27], and genome-scale metabolic modeling of community assembly has shown that higher community diversity is seen when each species occupies fewer niches [23]. This suggests that the conditions under which our model displays a positive diversity slope may be generally met in microbial communities, explaining our empirical observation of positive slopes in our experiment. Our model thus supports the idea that DBD can arise from cross-feeding interactions in our self-assembled communities.

Understanding the mechanisms that promote and sustain biodiversity in microbial communities is a central question in microbiome biology. Our paper addresses an important hypothesis: does diversity beget diversity in microbiomes? And more specifically, do more diverse microbiomes produce novel niches that recruit yet more taxa, further promoting diversity? While recent work had found a positive correlation between community diversity and focal diversity [22], this study focused on natural microbiome surveys where it is difficult to conclusively prove whether a positive correlation arises because of DBD or due to other environmental factors. By assembling microbial communities in a single nutrient– and so where coexistence is maintained via metabolic cross-feeding [26, 27] – we provide direct empirical evidence that diversity begets diversity in microbial community assembly and that it emerges via metabolic niche construction and cross-feeding interactions. Further, using consumer-resource models, we show that DBD is not inevitable in the presence of cross-feeding but that it depends on the species metabolic structure: DBD is more prevalent when secretions are both family-specific and sparse, and when species are resource specialists.

## Methods

### Community assembly experiment

Carbon source stock solutions (10X; 0.7 C-mol/L) were prepared at the start of the experiment, filter-sterilized (0.22um membrane filter), aliquoted in a 96-wellplate, and stored at −20C. Each transfer day, aliquots for each carbon source were thawed (single freeze/thaw cycle per aliquot) and diluted 10 fold into M9 minimal media. The final concentration of each carbon source is 0.07 C-mol/L. The 24 carbon sources used in the assembly experiment are listed in **Table S1**.

Soil samples from 8 different natural sites in New Haven and West Haven (CT, USA) were collected using the same sampling procedure as described in [10]. Once in the lab, each soil sample (5g) was suspended into 50ml of PBS 1x supplemented with 200ug/ml cycloheximide to inhibit eukaryotic growth, mixed, and left for 48h at room temperature. After the 48h, 40ul of each ‘source soil microbiome’ (inoculum) was inoculated into 500ul M9 minimal media supplemented with a single carbon source in 3 or 4 replicates (in a 96-well plate), and incubated at 30C for 48h. After the 48h-incubation period, each culture was mixed by pipetting, and then serially diluted (125x) into fresh medium. This growth/serial dilution cycle was repeated 10 times. Cycloheximide was added to the growth media in the first 2 transfers. OD620 was measured at the end of each incubation cycle. At transfer 10, the samples were centrifuged at 3500 rpm for 40min, and the pellets were stored at −80C until further processing.

### DNA extraction, sequencing, and taxonomy assignment

The pellets were thawed and the DNA was extracted using the DNeasy 96 Blood & Tissue kit (QIAGEN), including the pre-treatment for gram-positive bacteria, following the kit’s protocol. The 16S rRNA amplicon libraries were prepared and sequenced by Microbiome Insights, Vancouver, Canada. Briefly, the amplicon libraries were prepared using dual-barcoded primers targeting the 16S V4 region [33]. The PCR products were purified and normalized using the high-throughput SequalPrep 96-well Plate Kit, and the amplicon libraries were sequenced using the 300 bp paired-end kit v3.chemistry on a Illumina MiSeq platform. We used DADA2 to infer ESVs from each sample [34]. The demultiplexed fastq files (one forward and one reverse per sample) were used as input for DADA2 (version 1.6.0). The forward and reverse reads were truncated at position 240 and 160, respectively, and merged with a minimum overlap of 100bp (other parameters set to default values). Taxonomy assignment of each ESV was done using a naive Bayesian classifier method [35] trained on the Silva reference database (version 128). All samples were rarefied to the highest depth possible of 10779 reads.

### Testing for DBD and Null models

Following a similar approach to Madi et al (2020), here we tested for DBD by determining the slope of the regression line of the number of ESVs per focal family as a function of the number of non-focal families. ESVs with unassigned family taxonomy were excluded from the analysis, giving a total of 360 ESVs and 46 families for the equilibrium communities (i.e., at Transfer 10) across the 8 inocula, and a total of 369 ESVs and 66 families for the initial inocula (also referred to as Transfer 0) across 7 inocula (note that inoculum 2 was excluded here because of very low read count). The slopes are determined using a linear model (lm function in R) and p<=0.05 determines significance. A positive slope is consistent with DBD while a flat or negative slope is consistent with niche filling. Note that in our communities all ESVs belong to the same Order thus going deeper in evolutionary time is irrelevant.

To assess whether the observed slopes and R2 are significant (i.e., not likely to have occurred just by random chance), we used a null model that randomizes ESVs across samples. The ESVs randomization step was repeated 500 times, calculating the slope and R2 for each randomization. **Fig. S2** shows the distributions of the slopes and R2 obtained. The p-value for the slope (or R2) represents the fraction of ‘simulated’ (randomized) communities in which the slope (or R2) is greater than the slope (or R2) of the ‘real’ (observed) communities. A p-value lower or equal than 0.05 indicates that the observed values are significantly greater than expected by random chance.

### Consumer-Resource Model

Simulations were carried out using the Community Simulator package, an implementation of the MiCRM for Python [30, 36], with the additional features described in [32]. The model considers a pool of *S* species (consumers) and a set of *M* resources. Consumers are characterized by their uptake rates of the different resources, which are encoded in a consumer preference matrix **c** (where the element *c _iα_* corresponds to the uptake rate of resource *α* for species *i*). Resources are divided into classes and species into families such that every family displays preference (i.e., higher average uptake rates) for a different class of resources (in our simulations we consider the same number of resource classes and species families). The strength of this preference is modulated by a parameter *q* as described in [30], such that *q* = 0 corresponds to an ‘all generalists’ scenario (consumers show no preference for any resource class over another, thus there is no family structure in **c**) and *q* = 1 corresponds to an ‘extreme specialists’ scenario (consumers are unable to utilize resources outside their preferred class as the corresponding values in **c** are zero). Species secrete metabolites as byproducts when they utilize a resource. The amount of energy carried by resource *α* is given by a parameter *w_α_* set to 1 unit of energy per unit mass for every resource in all our simulations. The leakage fraction (*l*) determines what fraction of the energy consumed is secreted back to the environment in the form of byproducts. The remaining fraction of energy (1 – *l*) is allocated to growth, with the energy to growth rate conversion rate being denoted as *g* and set to 1 unit of energy^-1^ in all our simulations. Which byproducts are secreted, and in what proportions, is encoded in a 3-dimensional metabolic matrix **D** such that the element *D_iβα_* represents the fraction of energy flux in the form of resource *β* that is secreted by species *i* when it metabolizes resource *α*. Cell death was considered negligible in our timescales, therefore we set the death rate (*m*) to zero in all our simulations.

In each set of simulations, communities were assembled by randomly sampling species from the pool, providing them with a primary resource (1000 units of mass at *t* = 0), allowing them to grow to equilibrium, diluting the final abundances of all species and resources, replenishing the primary resource, and repeating the process for 20 cycles. At that point, the composition of the communities at the end of a cycle did not undergo significant further changes. The dynamics of the species and resources in the community between consecutive dilutions was simulated by numerically integrating the following equations as described in [30]:

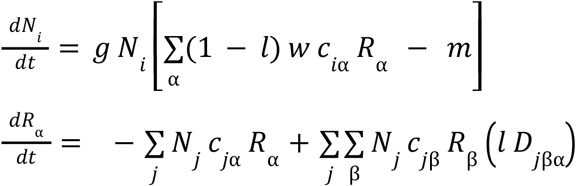

where *N_i_* represents the abundance of species *i* and *R_α_* represents the abundance of resource *α*.

We ran a total of 100 sets of simulations, assembling 100 communities in each. For each set, model parameters were randomized by sampling uniformly within the following ranges: total number of families in the pool between 3 and 8; total number of species per family in the pool between 150 and 300; number of species sampled from the pool to initialize the communities between 20 and 100 times the number of families; leakage fraction between 0.5 and 0.8; consumers preference strength *q* between 0 and 1. Matrices **c** and **D** were also resampled for each set of simulations. The elements of **c** were sampled from a Gamma distribution, while each column of **D** was sampled from a Dirichlet distribution such that the condition 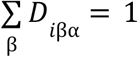 (which ensures conservation of energy, i.e., that there is not more energy being secreted than uptaken by any consumer) is satisfied for every species *i* and consumed resource *α*. The elements of **D** were sampled such that 45% of the secreted energy flux went into resources of the same class as the consumed one, the default for the Community Simulator package [30]. We also made it so secretions could be the same across families (*D_iβα_* = *D_jβα_* for any two species *i* and *j* regardless of whether they belong to the same or different families, and for any combination of consumed/secreted resources *α* and *β*) or allowed for family-specific secretions (*D_iβα_* = *D_jβα_* if species *i* and *j* belong to the same family, but may be different if *i* and *j* belong to different families). The sparsity of the matrix **D** determines how distributed secretions are across resources. This behavior is controlled in the model by a sparsity parameter as described in [31], such that when it is small (≪1) secretions tend to be distributed across all resources, and when it is large (≫1) they tend to be clustered across few resources. In each set of simulations, this scarcity parameter was sampled between 0.1 and 10 with uniform probability in the log scale.

We considered that a simulation regime corresponded to a ‘strong specialists’ scenario if the preference strength *q* was above 0.5, and to a ‘weak specialists’ (or ‘generalists’) scenario otherwise. Secretions were considered ‘sparse’ if the sparsity parameter used to construct the matrix **D**was greater than 4, and ‘distributed’ otherwise. Finally, we considered that there was a ‘common core metabolism’ if *D_iβα_* was equal to *D_jβα_* for every pair of species *i* and *j*, and a ‘family-specific’ metabolism if they could be different whenever *i* and *j* belonged to different families as described above.

## Acknowledgments

We would like to thank Josh Goldford for helpful discussions. This work was supported by a Packard Foundation Fellowship to A.S. and by the National Institutes of Health through grant 1R35 GM133467-01 to A.S.

## Supplementary Text

### A null model of community assembly

We consider a pool of an arbitrary number *S_tot_* of species divided into an arbitrary number *F* of families, one of which is the focal (**Fig. S1A**). We denote the number of species that belong to family *i* as *S_i_*. (such that 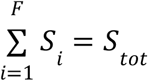) and the population size (number of individuals) of species *j* from family *i* as *N_ij_* (with *j* = 1, 2,…, *S_i_*). The total number of individuals of family *i*, denoted as *N_i_,* is thus 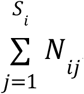. The total population size of the pool, denoted as 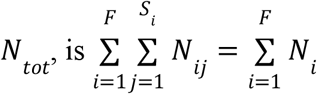. We examine whether assembling a random community by sampling *n* individuals from the pool is expected to result in a positive diversity slope (**Fig. S1A**).

The process of sampling individuals from the pool one at a time can be seen as a random walk of changing probabilities in the plane defined by the number of non-focal families and the number of focal species in the sample (**Fig. S1B**). Sequentially drawing individuals is equivalent to taking steps in this walk, either ‘to the right’ if the drawn individual belongs to a family not sampled previously, or ‘up’ if the drawn individual belongs to the focal family and was not sampled previously. Naturally, it is not possible to take steps ‘down’ or ‘to the left’ if we are not allowing for the loss of individuals already in the sample; however, it is possible to ‘stay in the same position’ if the individual drawn is of a focal species that is already present in the sample, or if it belongs to a non-focal family that is already represented in the sample. As the drawing process advances, the probabilities of sampling new families or focal species decreases (**Fig. S1B**, inset). Specifically, at the stage where the sample contains *m* < *n* individuals, the average probabilities *p_up_* and *p_right_* of taking further steps ‘up’ or ‘right’ can be respectively approximated by:

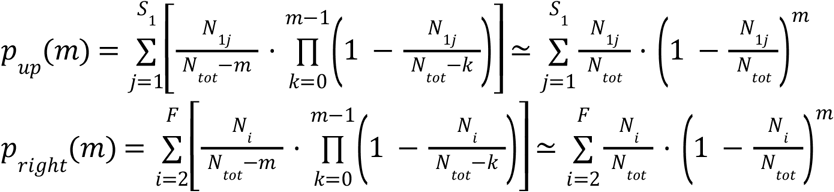

where we have assumed, without loss of generality, that the first family is the focal. We have also assumed that population sizes are sufficiently large (*≫ m*) for the probabilities to be well approximated by those of a sampling with replacement. These probabilities approach zero as the sample size increases, and ultimately the random walk ends when all non-focal families and focal species have been sampled.

Because the process only allows for net ‘up and right’ movement, when one considers a set of random walks (that is, a set of communities randomly assembled in this fashion) a positive diversity slope is expected to emerge. How *p_up_* and *P_right_* vary with *m* depends on the distribution of species population sizes in the pool, i.e., on the specific values of *N_ij_*. Typically, *P_right_* will start off higher than *p_up_* if the focal family is not overwhelmingly represented in the pool, that is, if the population size of the focal family does not exceed that of all other families combined:

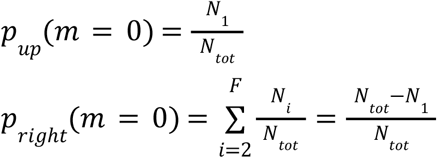

If the probability of a ‘step right’ is much larger than that of a ‘step up’ around *m* ~ 0, we expect to see only a weak diversity slope for very small sample sizes, since any set of random walks will tend to exhibit notably larger variation in the horizontal than in the vertical direction. As *m* slowly increases, *p_up_* and *p_right_* decay at a rate that is again determined by the distribution of population sizes in the pool. For very small values of *m,* the probabilities change approximately as:

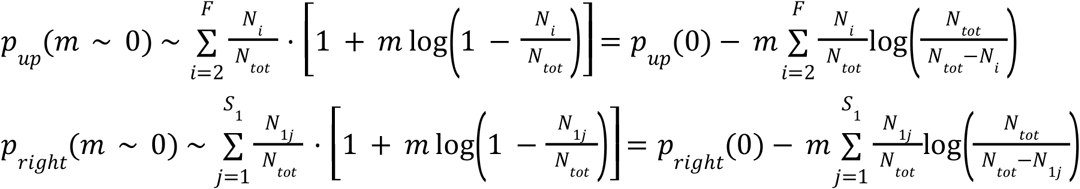

As probabilities decay and get closer to one another, a stronger positive diversity slope could emerge because ‘diagonal movement’ becomes more favored in the random walk. Finally, for very large sample sizes both *p_up_* and *P_right_* approach zero, thus in that regime we again expect a weak diversity slope due to the lack of significant variation in both axes.

To test our intuition regarding the emergence of a positive diversity slope in this null model, we first constructed a pool of species divided into 50 families, with 30 to 60 species in each of them. Within each family, species population sizes were between 10^4^ and 10^8^ individuals and log-normally distributed. We considered three regimes: small sample sizes (5-25 individuals), intermediate sample sizes (500-2,500 individuals) and large sample sizes (50,000-250,000 individuals). In each regime, we assembled 100 communities by sampling a random number of individuals *n* from the pool, with *n* uniformly distributed within the ranges specified before. Our choice of intervals makes it so the relative variation in sample sizes within each regime is roughly maintained (coefficient of variation ~ 0.4 in all three regimes). We then quantified the number of non-focal families and the number of focal species of each community of every community. **Fig. S1C** shows that, in line with our previous reasoning, a strong positive diversity slope could emerge at intermediate sample sizes.

## Supplementary Figures and Tables

**Fig. S1.**
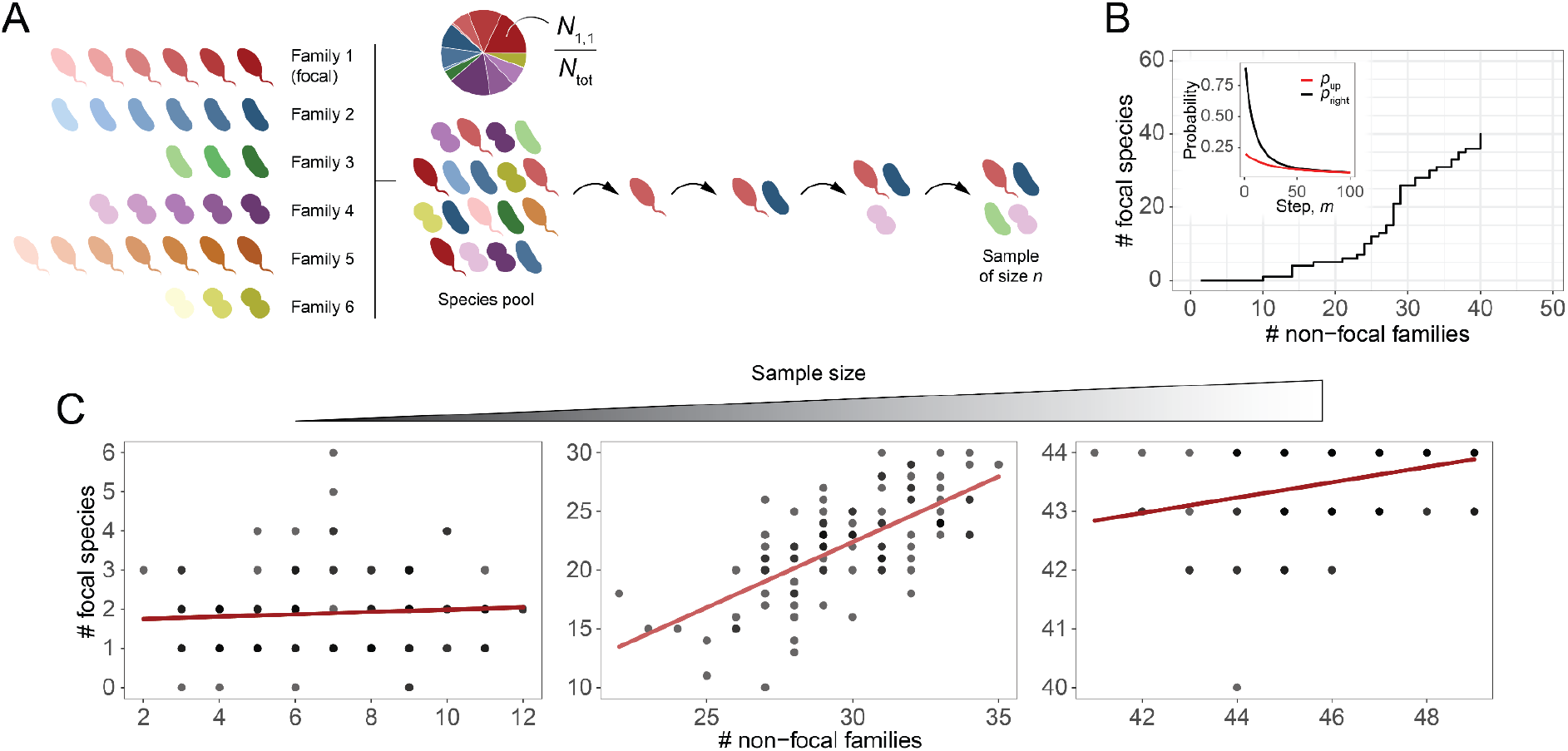
A positive diversity slope can emerge from the random sampling of a species pool in the absence of species interactions. (**A**) We consider a pool of an arbitrary number of species divided into an arbitrary number of families, out of which the first one is the focal. The pie plot represents the relative abundances of each species in the pool. We examine whether assembling a random community by sequentially drawing *n* species from the pool is expected to give a positive diversity slope. (**B**) The sampling process can be seen as a random walk of changing probabilities. Drawing species from the pool one at a time is equivalent to taking steps in this walk, either ‘to the right’ if the drawn species belongs to a family not sampled previously, or ‘up’ if the drawn species belongs to the focal family and was not sampled previously. An example walk is represented here. The inset shows how the average probabilities of taking a step ‘up’ or ‘to the right’ change as one keeps drawing more species from the pool, that is, taking more steps. (**C**) We simulated 100 random draws from a species pool analogous to the one in panel A, with species abundances log-normally distributed in the pool. We tested three different regimes: small, intermediate and large sample sizes (*n* =5 to 25, 500 to 2,500, or 50,000 to 250,000 individuals, respectively). We quantified the number of non-focal families and of focal species for each sample. A strong positive diversity slope emerged at intermediate sample sizes.

**Fig. S2.**
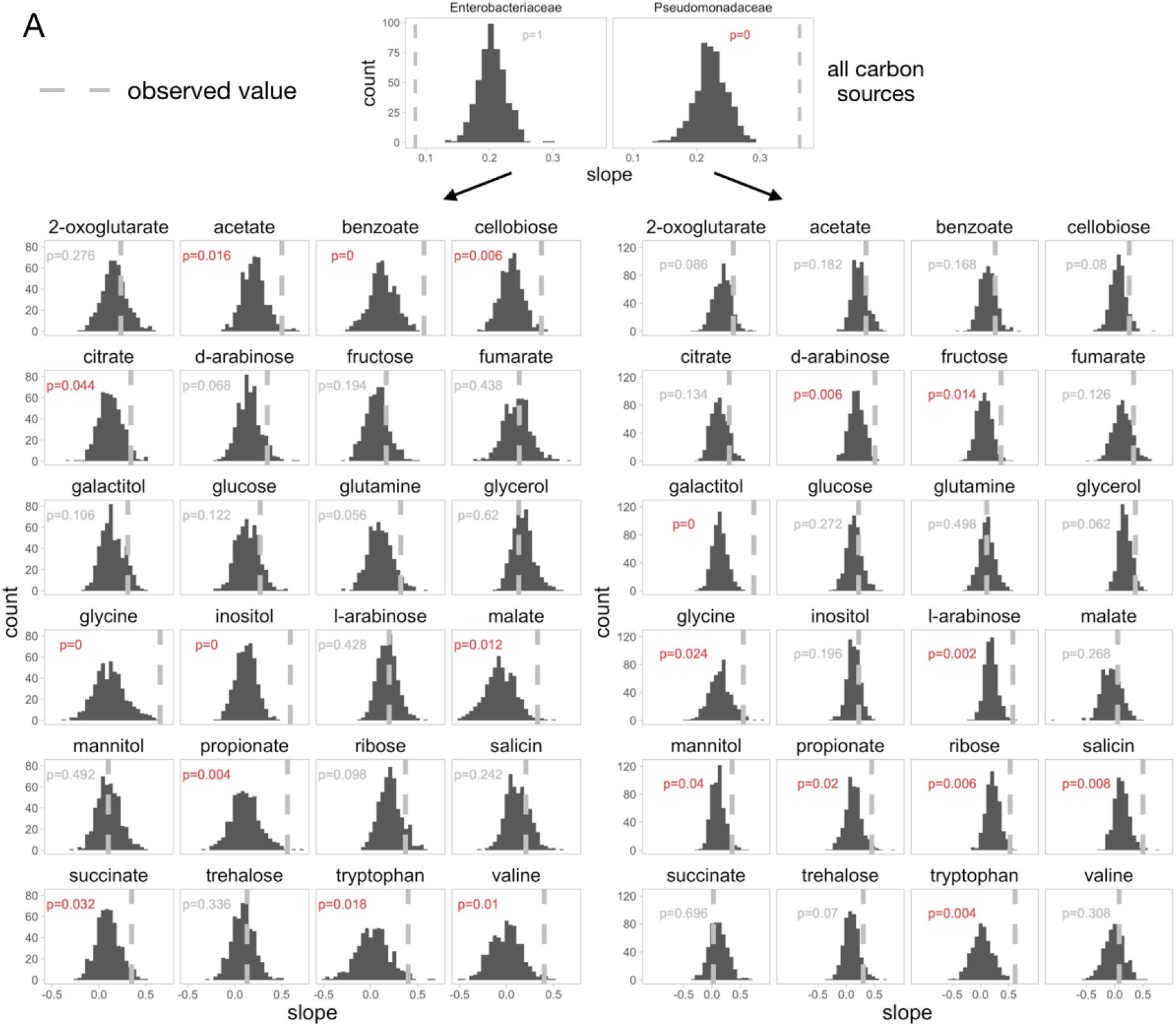
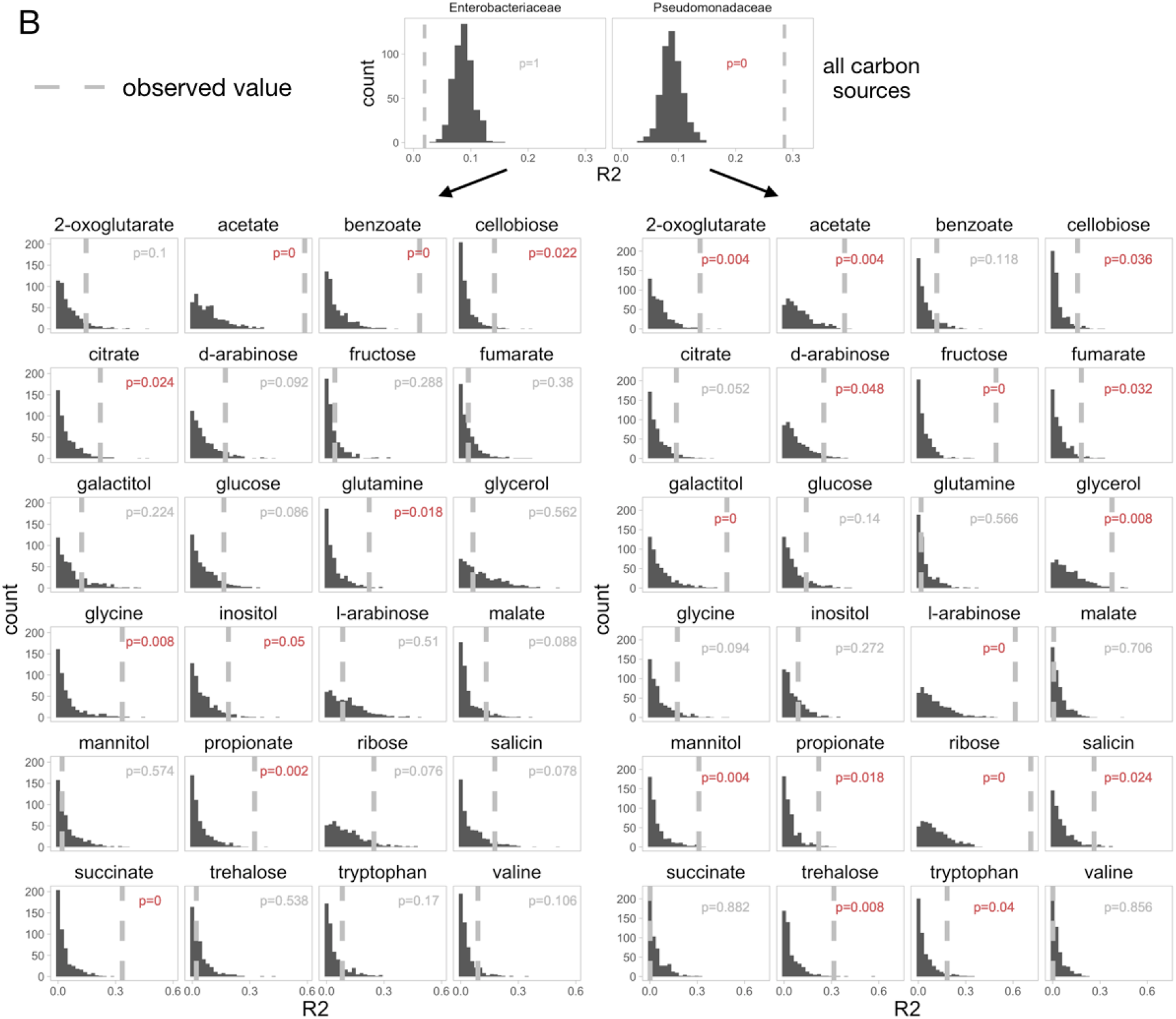
Histograms showing the distributions of the diversity slopes (A) and R2 (B) of the randomized (simulated) communities for Enterobacteriaceae and Pseudomonadaceae across all carbon sources or each carbon source separately. N=500 randomizations. Dashed lines indicate the value of the observed (real) communities. The p-values represent the fraction of randomized communities in which the slope (or R2) is greater than the slope (or R2) of the observed communities. p<= 0.05 indicates that the observed values are significantly greater than expected by random chance, and are shown in red.

**Fig. S3.**
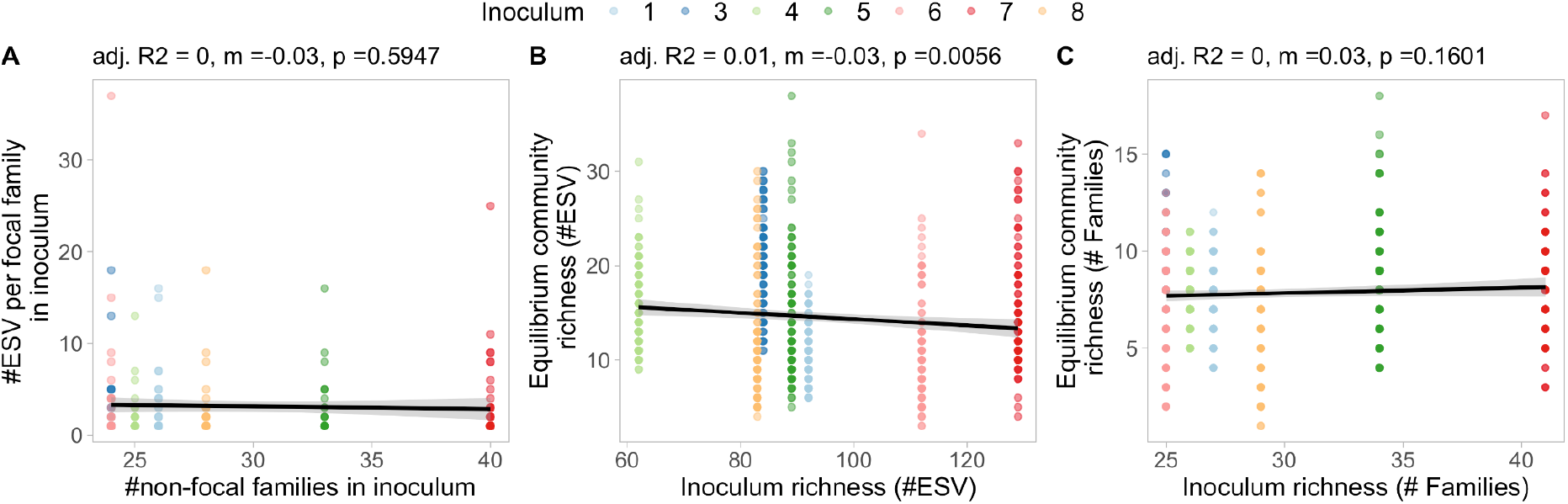
Community richness is not positively correlated with inoculum richness. (**A**) Shown is the number of ESVs per focal family in the initial inoculum (T0) as a function of the number of non-focal families at T0 (N=207). (**B**, **C**) Shown is the number of ESVs (or families) in the communities at equilibrium (T10) as a function of the number of ESVs (or families) in the initial inoculum (T0), colored by inoculum (N=649). Values from linear regression are shown above each plot (m: slope, p: p-value). All communities were rarefied to the same sequencing depth of 10779 reads. Inoculum 2 was excluded from the analysis because of very low read count in T0.

**Table S1.** Carbon sources used in this study

| **Name** | **Brand** | **SKU** |
| --- | --- | --- |
| D-Glucose | VWR | 0188-500 |
| D-Mannitol | Sigma | M4125-100 |
| D-Cellobiose | Sigma | 22150-10G |
| D-Trehalose Dihydrate | Sigma | 90210-50G |
| D-Fructose | Acros Organics | 161355000 |
| D-Ribose | Acros Organics | AC132361000 |
| L-Arabinose | DOT Scientific | DSA51000-500 |
| D-Arabinose | Sigma | 10850-25G |
| D-Salicin | Alfa Aesar | A16524 |
| Glycerol (80%, w/v) | Teknova | G8797 |
| Dulcitol | Sigma | D0256-100G |
| Inositol (myo-inositol) | Sigma | 57570-25G |
| Sodium citrate dihydrate | Fisher | S279-3 |
| 2-Ketoglutaric acid, disodium salt, dihydrate | Acros Organics | 271250500 |
| Sodium Succinate hexahydrate | Alfa Aesar | 419A3 |
| Sodium hydrogen fumarate | Alfa Aesar | B24683 |
| Sodium acetate tri-hydrate | Sigma | S7670-250G |
| Sodium propionate | Alfa Aesar | A1744036 |
| Sodium benzoate | Alfa Aesar | A15946 |
| L-Glutamine 200mM (29.23 mg/mL) | Sigma | G7513-100ML |
| L-Valine | Sigma | 94619-25G |
| Glycine | Sigma | G7126-100G |
| L-Tryptophan | Sigma | TO254-25G |
| L-(–)-Malic acid sodium salt | Sigma | M1125-5G |

